# Gray fox (Urocyon cinereoargenteus) survival in southern Illinois, USA

**DOI:** 10.64898/2026.08.20.746046

**Authors:** Nadine A. Pershyn, Clayton K. Nielsen, Guillaume Bastille-Rousseau

## Abstract

Gray fox (*Urocyon cinereoargenteus*) populations in the Midwestern USA have suffered precipitous declines in recent decades, yet they are relatively understudied. However, understanding survival and cause-specific mortality is vital for declining populations and the limited existing survival studies have been performed outside of the Midwest. We equipped 13 gray foxes in southern Illinois with GPS radio collars to investigate their survival and cause-specific mortality. We calculated the Kaplan-Meier 6- and 12-month survival rates to be 0.79 (95% CI: 0.57-1.0) and 0.53 (95% CI: 0.27-1.0), respectively. We recorded 4 mortalities: 1 disease, 1 gunshot, and 2 unknown causes. While our study has a small sample size, it contributes key information on a data-deficient mesocarnivore suffering from a population decline driven by undefined causes. We recommend further research into the survival and mortality of this elusive mesocarnivore.

## INTRODUCTION

Gray foxes (*Urocyon cinereoargenteus*) have a wide distribution across Central and North America (Fritzell and Haroldson 1982), however, populations have declined precipitously in the Midwest during the last 40 years (Cooper *et al*. 2012; Lesmeister *et al*. 2015; Bauder *et al*. 2020; Larreur *et al*. 2025). Despite their status as a furbearing mammal they are an understudied mesocarnivore; less is known about gray foxes than most other furbearers (Sillero *et al*. 2004).

There are limited studies on gray fox distribution and population status (Cooper *et al*. 2012; Lesmeister *et al*. 2015; Allen *et al*. 2021), and fewer demographic studies, especially as it relates to recruitment, sex- and age-specific survival, and fecundity (Allen *et al*. 2021).

Gray foxes are often associated with deciduous and/or coniferous forests interspersed with some fields and scrubby woodlands (Hall 1981). Gray foxes are the most omnivorous of all North American fox species, and their diets include insects, birds, fruits and nuts, and sometimes carrion, however during the winter they consume primarily rabbits (*Sylvilagus* spp.) and rodents (Fritzell and Haroldson 1982). Across their range, gray foxes are listed as ‘Least Concern’ by the International Union for Conservation of Nature (Roemer *et al*. 2016). However, the Illinois Department of Natural Resources (IDNR) currently designates gray foxes as a ‘watch list’ species, which indicates they have poorly known distributions, trends, and habitat requirements (IDNR 2005). Annual harvests in Illinois during 1976–1989 were ~3,200 individuals but declined to <40 by 2015 (Bauder *et al*. 2020; Williams *et al*. 2020). Gray foxes experienced much higher extinction rates than colonization rates in southern Illinois between 2010 and 2024 (Lesmeister *et al*. 2015; Larreur *et al*. 2025), which is indicative of a declining population.

However, there have been relatively minimal changes to land cover composition over the past decades (Walk *et al*. 2010), suggesting that habitat change is not propelling this decline. This is further supported by evidence that current gray fox occupancy in southern Illinois is inconsistent with adapting to landcover change alone (Larreur *et al*. 2025). Rather, coyotes may be an emerging problem for gray fox, as they are intraguild predators that can decrease gray fox occupancy and abundance (Lesmeister *et al*. 2015; Egan *et al*. 2021). Additionally, disease, particularly canine distemper can have detrimental impacts on gray foxes (Davidson *et al*. 1992).

Survival and mortality are major contributors to population status and viability (Morris and Doak 2004). Therefore, when populations are declining, it is essential to quantify survival rates as they provide a standardized measure that can be compared between populations.

Additionally, identifying the main causes of death is critical if managers intend to combat the decline. Research investigating gray fox survival in the Midwest is imperative to assess drivers of the observed population decline. Previous studies of gray fox outside the Midwest reported mean annual survival rates between 0.58 and 0.69 (Chamberlain 1999; Weston and Brisbin 2003; Farias *et al*. 2005; Temple *et al*. 2010). The highest rate in the literature is based on an age structured Krebs model for gray foxes in the Savannah River Site nuclear production facility (Weston and Brisbin 2003). However, the remaining survival rates based on radio-collared individuals are all very similar, with three estimates falling between 0.58-0.61 (Chamberlain 1999; Farias *et al*. 2005; Temple *et al*. 2010). One previous gray fox study in the Midwest did not estimate survival rate due to too small of a sample size, only monitoring 7 foxes (Cooper 2008). Studies reported that causes of gray fox mortality included predation, human caused (i.e., harvest and vehicle collisions), and disease (i.e., canine distemper, canine hepatitis, and rabies) (Weston and Brisbin 2003; Chamberlain and Leopold 2005; Farias *et al*. 2005; Cooper 2008; Temple *et al*. 2010).

We quantified survival of gray foxes in southern Illinois and investigated the causes of mortality in this region. We hypothesized that survival would be lower than reported survival of gray fox outside of the Midwest, and that coyote predation and disease would be leading causes of mortality.

## METHODS

### Study Area

We conducted this study in Gallatin, Hardin, Pope Saline, and Johnson counties of southern Illinois, USA (Figure 1), which had a human population density of 10.5 persons/km^2^ (United States Census Bureau 2022). Elevation ranged from 91 to 325 m (Netstate 2023). Annual mean air temperatures in southern Illinois were 5.4 ± 1.4 °C and annual mean precipitation was 36.3 ± 10.7 cm (National Oceanic and Atmospheric Administration [NOAA] 2021). The study area was 910 km^2^ and was comprised of 55% deciduous forest, 13% grassland, 9% agriculture, 9% mixed forest, 5% coniferous forest, 4% developed, and <1% each of herbaceous shrub, wetlands, and barren ground (Dewitz 2023). Road density was 1.4 km/km^2^ and stream density was 1.1 km/km^2^.

**Figure 1.**
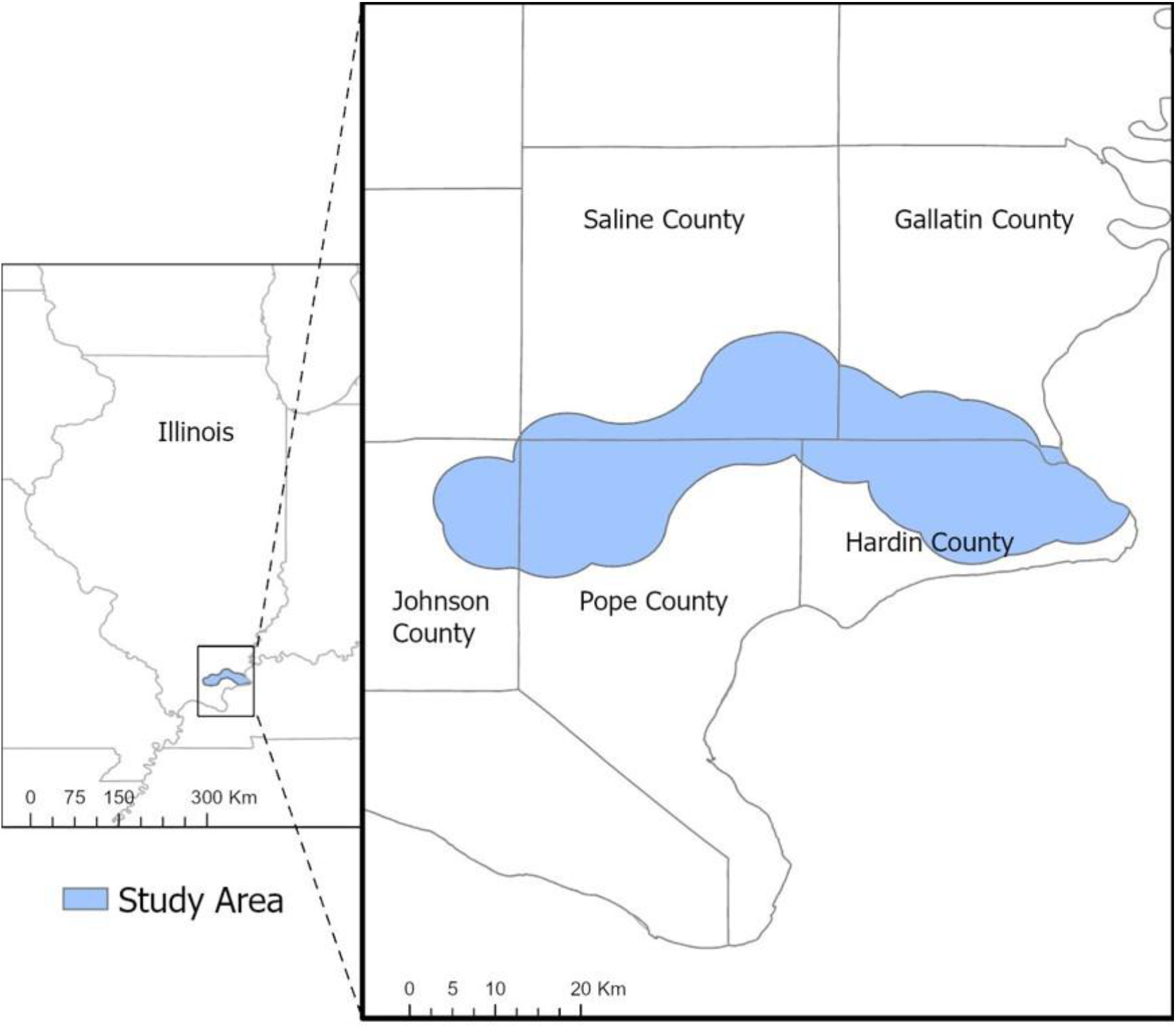
Study area map for a gray fox survival study in southern Illinois, USA, 2022-2025.

### Capture and Handling

Capture locations were chosen based on 2022-2024 gray fox detections and occupancy models (Larreur *et al*. 2025). Scouting cameras were placed near areas where camera traps captured an image of a gray fox, or areas that occupancy modeling suggested had relatively high probabilities of gray fox occupancy. Traps were deployed in areas where camera traps confirmed the presence of gray fox. We used padded number 1.5 Soft-catch (Oneida Victors’, Cleveland, OH, USA) foothold traps to capture gray foxes between November-March for three consecutive field seasons during 2022-2025 with commercial baits and gland lures used as attractants (Minnesota Trapline Products, Pennock, MN, USA). Traps were primarily placed on Shawnee National Forest land, with some on private land with permission from landowners. Traps were placed >100 m from any inhabited dwellings, >2 m off established trails, and/or directly on game trails or at the base of trees and logs.

Gray foxes were handled under chemical immobilization using BAM (butorphanol, azaperone, medetomidine). Anesthetized foxes were blindfolded and fitted with a muzzle for additional safety. We took blood samples, affixed ear tags, and recorded weight, sex, and morphological measurements. A 10 mm x 1.41 mm microchip (AVID; Friendchip-Mini) was injected under the skin between the shoulder blades to help identify re-captured individuals. Individuals were classified as adults (>1 yr) or juveniles based on body mass and condition of dentition. Gray foxes were fitted with LiteTrack Iridium-150 GPS collars (LotekWireless, Newmarket, Ontario, Canada) programmed to record locations every 2 hr and with a release mechanism to drop off one year after activation. Capture and handling protocols were reviewed and approved by the Institutional Animal Care and Use Committee (IACUC) at Southern Illinois University Carbondale (#22-020) and in accordance with guidelines endorsed by the American Society of Mammologists (Sikes *et al*. 2011).

### Survival Rates

Deceased animals were found and cause of mortality determined following necropsy, classified in the following categories: predation, disease, anthropogenic sources, unknown. We estimated 6-month and 12-month (annual) survival rates using the Kaplan-Meier (KM) method (Pollock *et al*. 1989). Due to low sample size, we combined data across 3 years by calculating survival as days since December 12^th^, which was the earliest capture date for any fox. For the 6-month survival estimate, we used June 12^th^ as the end date of the analysis. Using the ‘survival’ package (Therneau 2023) in R version 4.3.1 (R Core Team 2023) we calculated staggered entry survival and included dates of right-censoring (i.e., collar failure, survival at end of the analysis).

## RESULTS

In 7,375 trap nights we captured 14 gray foxes (5 ad F, 6 ad M, 1 juv F, 2 juv M; Table 1) between 2023-2025. We collected a total of 15,764 GPS locations, and locations per individual ranged from 124 to 3,489 (mean 1,204 ± 1,027 (SD); Table 1). Excluding mortalities and a slipped collar, collars were active between 44 to 362 days (mean 164 ± 108 (SD); Table 1) until they stopped reporting data, or in one case the timed release device (i.e., collar drop-off mechanism) activated early. We recorded 5 mortalities: 1 capture myopathy (which was censored from the study), 1 disease, 1 gunshot, and 2 instances where the carcass was too decomposed/scavenged to determine cause of death (i.e., unknown mortality cause). The majority of foxes were right-censored from the survival analysis due to collars not reporting data for a complete year. Based on 2 mortality events, we calculated the 6-month KM survival to be 0.79 (n =13, 95% CI: 0.57-1.0; Figure 2a). Based on 4 mortality events, we calculated the KM annual survival for gray foxes in the study area to be 0.53 (n = 13, 95% CI: 0.27-1.0; Figure 2b).

**Table 1.**
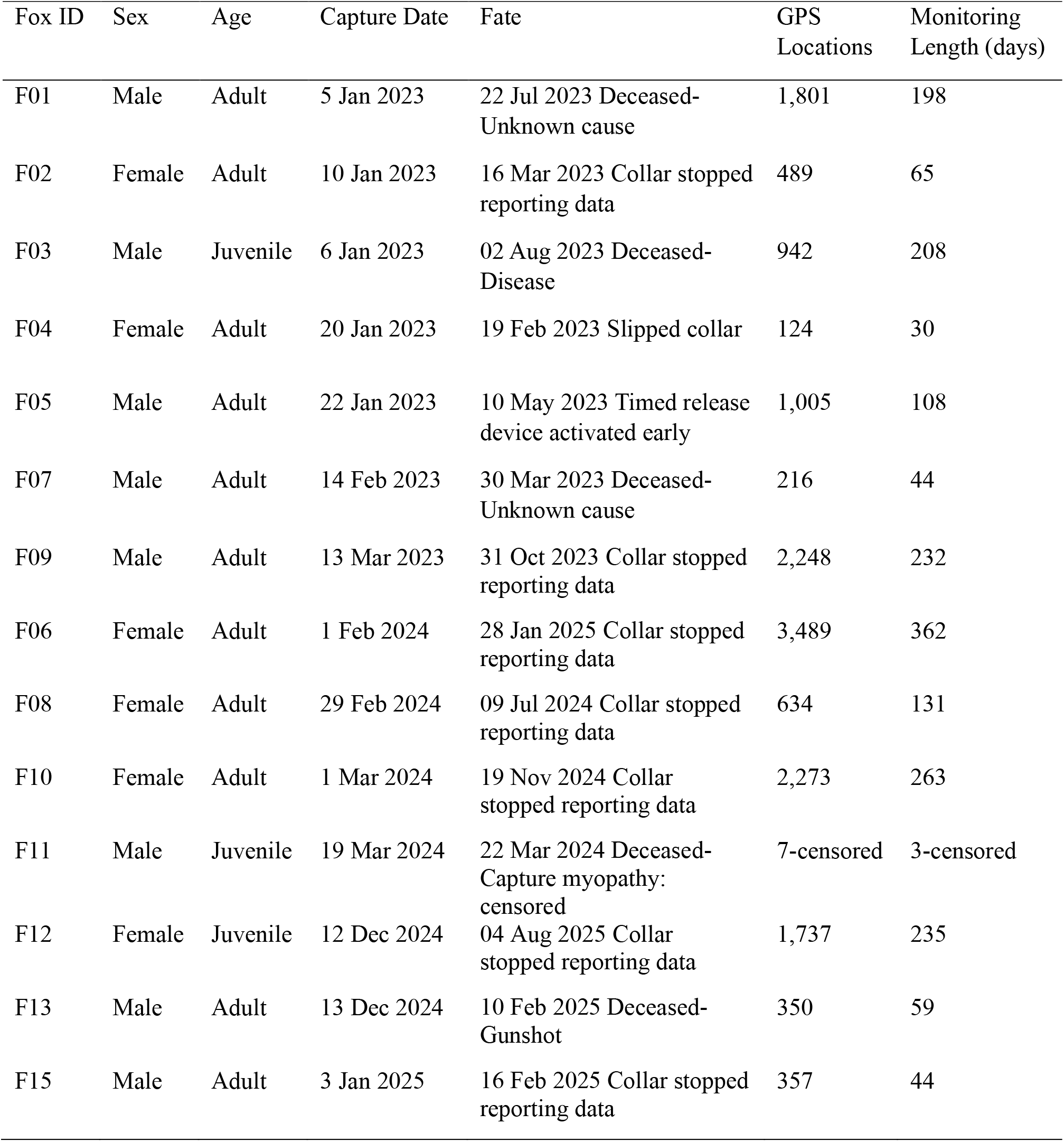
Gray foxes captured in southern Illinois, USA, 2022-2025. 14 individuals were captured (ID F14 was not assigned) and 13 yielded useful data (F11 died due to capture myopathy and was censored from the study).

| Fox ID | Sex | Age | Capture Date | Fate | GPS Locations | Monitoring Length (days) |
| --- | --- | --- | --- | --- | --- | --- |
| F01 | Male | Adult | 5 Jan 2023 | 22 Jul 2023 Deceased-Unknown cause | 1,801 | 198 |
| F02 | Female | Adult | 10 Jan 2023 | 16 Mar 2023 Collar stopped reporting data | 489 | 65 |
| F03 | Male | Juvenile | 6 Jan 2023 | 02 Aug 2023 Deceased-Disease | 942 | 208 |
| F04 | Female | Adult | 20 Jan 2023 | 19 Feb 2023 Slipped collar | 124 | 30 |
| F05 | Male | Adult | 22 Jan 2023 | 10 May 2023 Timed release device activated early | 1,005 | 108 |
| F07 | Male | Adult | 14 Feb 2023 | 30 Mar 2023 Deceased-Unknown cause | 216 | 44 |
| F09 | Male | Adult | 13 Mar 2023 | 31 Oct 2023 Collar stopped reporting data | 2,248 | 232 |
| F06 | Female | Adult | 1 Feb 2024 | 28 Jan 2025 Collar stopped reporting data | 3,489 | 362 |
| F08 | Female | Adult | 29 Feb 2024 | 09 Jul 2024 Collar stopped reporting data | 634 | 131 |
| F10 | Female | Adult | 1 Mar 2024 | 19 Nov 2024 Collar stopped reporting data | 2,273 | 263 |
| F11 | Male | Juvenile | 19 Mar 2024 | 22 Mar 2024 Deceased-Capture myopathy: censored | 7-censored | 3-censored |
| F12 | Female | Juvenile | 12 Dec 2024 | 04 Aug 2025 Collar stopped reporting data | 1,737 | 235 |
| F13 | Male | Adult | 13 Dec 2024 | 10 Feb 2025 Deceased-Gunshot | 350 | 59 |
| F15 | Male | Adult | 3 Jan 2025 | 16 Feb 2025 Collar stopped reporting data | 357 | 44 |

**Figure 2.**
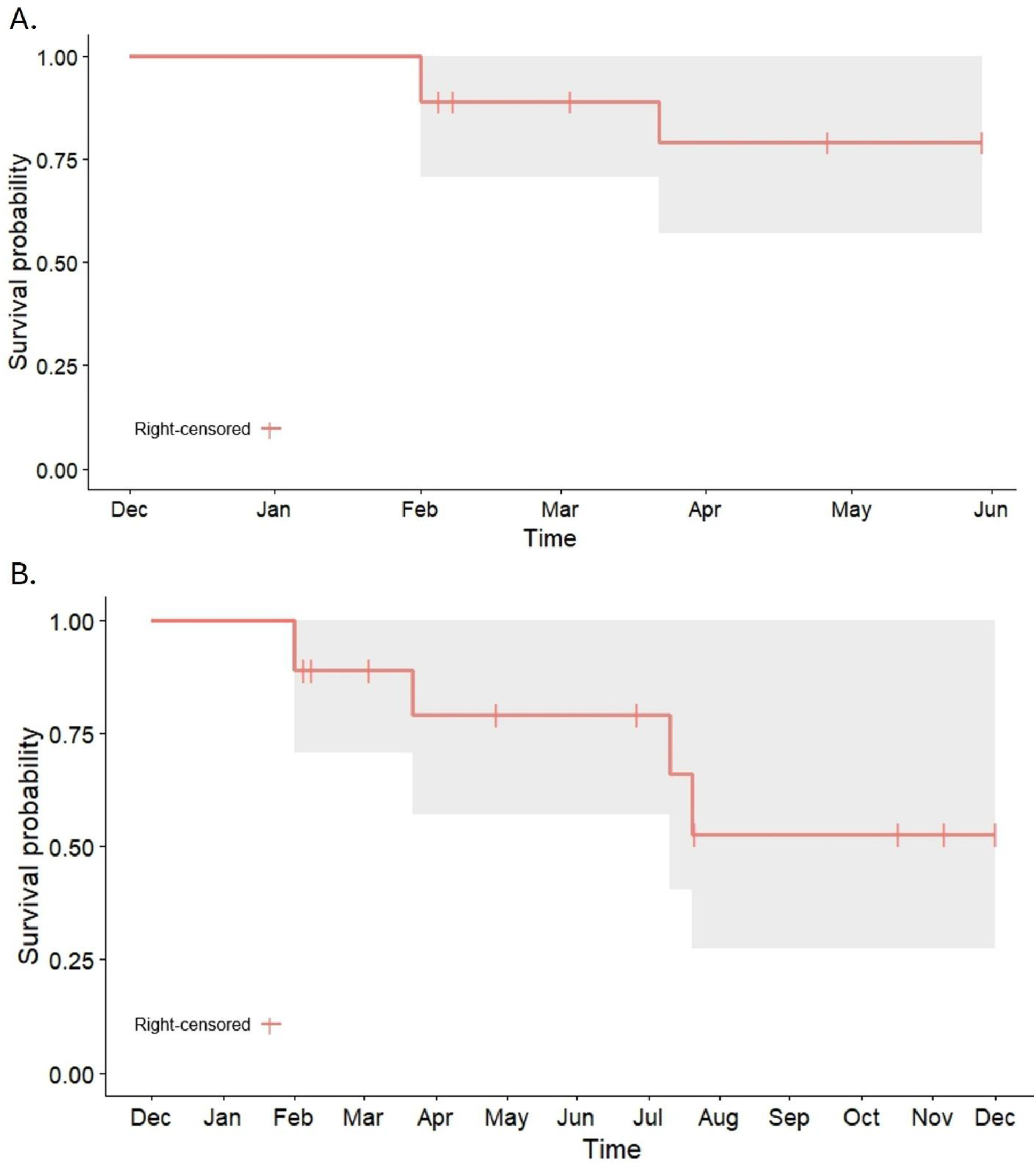
Kaplan-Meier (A) 6-month and (B) annual survival curves of n = 13 gray fox in southern Illinois, USA, 2022-2025. Drops in the main horizontal line indicate mortalities and vertical dashes represent right-censored individuals. 95% confidence intervals are indicated by gray shading.

## DISCUSSION

We calculated an annual survival rate lower than any previously published gray fox survival rate, which supported our hypothesis, however the confidence intervals encompassed all other published estimates, so our estimate was not significantly different. The lowest survival rate found in the literature was 0.58 (95% CI: 0.39-0.85) and was calculated from gray foxes in southern California in 1997-1999 (Farias *et al*. 2005). While our mean survival rate of 0.53 (95% CI: 0.27-1.0) fell below that, our uncertainty was extraordinarily large due to our sample size constraints. As a result of the low sample size, we were unable to investigate sex- or age-specific survival, or to account for annual differences. Other studies have documented similar survival across years and between sexes (Chamberlain 1999; Farias *et al*. 2005; Temple *et al*. 2010), while adult survival is typically higher than juvenile survival (Farias *et al*. 2005).

We experienced difficulties at multiple stages that resulted in low statistical power for our analysis. First, most likely as a result of population declines, we struggled to locate foxes to trap; we even noticed a decrease in gray foxes over the course of the three trapping seasons.

Additionally, an overlapping study conducted during 2022-2024 with extensive trail camera grids recorded no gray fox sightings in 2024 (Larreur *et al*. 2025). Second, the collars deployed on the foxes we caught did not report data for long enough; only one came close to lasting the year anticipated during project development. The majority stopped reporting data much sooner, with one failing a mere 6 weeks after deployment. The lack of long-term monitoring was detrimental to the survival analysis, as individuals were censored before a mortality event could occur or they could survive a full year. Finally, when the few mortality events occurred, we were not always able to retrieve the carcasses in time to determine the cause of death. In one instance, the mortality alert did not send until two days after a landowner saw the dead fox on their property.

This could be due to a delay in the collar sending the alert, or due to scavengers moving the body and keeping the collar active. Additionally, the extreme heat and humidity of late spring/summer contributed to fast decomposition of mortalities during those times, which created further difficulties. Therefore, half of our natural mortalities were unknown causes, which greatly impedes the ability to identify forces potentially driving the decline.

We identified two mortality sources, disease and gunshot, but did not identify any mortality due to predation, which contradicted our hypothesis. A previous gray fox study in southern Illinois recorded 4 non-capture related mortalities, 2 (50%) from coyote predation, 1 (25%) from a ringworm infection, and 1 (25%) unknown cause (Cooper 2008). Disease, especially canine distemper, is a large concern for gray foxes (Davidson *et al*. 1992). However, we were unable to identify the specific disease responsible for the mortality. At the time of the gunshot mortality, the gray fox hunting/trapping season was in effect, although this individual was not thereafter collected by the hunter/trapper but recovered by researchers using the mortality sensor. After the conclusion of the final field season in 2025, the Illinois gray fox hunting and trapping season was closed indefinitely to remove additional pressure and additive mortality from harvest (IDNR 2025). No radio-collared foxes suffered from vehicle mortality, which can be a leading cause of mortality for gray foxes (Temple *et al*. 2010). However, one non-collared gray fox was identified as roadkill in the study area by the Illinois Department of Natural Resources during the study (A. Phillips, personal communication, October 17, 2023). While predation by coyotes or occasionally bobcats is often identified as a mortality source for gray foxes (Farias *et al*. 2005; Cooper 2008), we could not confirm any predation mortality.

Our study contributes to the limited literature on gray fox survival and, despite the high uncertainty in our estimates, provides critical data to an understudied mesocarnivore during population decline. Further research is necessary to gain a better understanding of gray fox survival and cause-specific mortality in the Midwest. While trapping foxes when populations are very low is difficult, larger sample sizes are necessary to draw population-scale conclusions and all efforts should be made to increase sample size including partnering with local trappers when feasible. Furthermore, improved GPS collar performance is crucial to the long-term monitoring necessary in survival studies. Gray foxes are particularly difficult to track with GPS collars as they often utilize dens located in hollow logs or underground (Nicholson *et al*. 1985), which makes it difficult to acquire GPS fixes, send data via satellite, and overall optimize battery use on GPS collars. Researchers should carefully consider the collar best suited to their project, whether that be iridium, store-onboard, ultra-high frequency (UHF) download, or some other collar.

Iridium relays GPS data automatically but uses more battery to do so; store-onboard collars can have increased battery life due to not sending data via satellites, however if the collars are not recovered all the data is lost; UHF download requires routinely relocating the animal and getting close enough to download the points via a UHF signal, which can be difficult or impossible if the animal disperses and is lost (Dore *et al*. 2020). For a survival study, we recommend prioritizing battery longevity to increase the likelihood of collecting sufficient mortality data, which could mean programming as few iridium connections as feasible and/or longer time between fixes.

Beyond having the data of when foxes die, understanding the specific mortality sources for gray foxes is essential to address the decline across the Midwest. Therefore, data should be collected in a timely manner, such that expedient investigation of mortalities can contribute to identifying cause of death. Additional research into gray fox survival and cause-specific mortality is critical,, especially as limited evidence outside the Midwest suggest they might be facing a more widespread threat (Rodgers 2026).

## LITERATURE CITED

Allen M, Avrin A, Farmer M, Whipple L, Cervantes A, Bauder J. 2021. Limitations of current knowledge about the ecology of Grey Foxes hamper conservation efforts. Journal of Threatened Taxa 13:19079–19092. 10.11609/jott.7102.13.8.19079-19092

Bauder JM, Allen ML, Ahlers AA, Benson TJ, Miller CA, Stodola KW. 2020. Identifying and Controlling for Variation in Canid Harvest Data. The Journal of Wildlife Management 84(7):1234–1245. 10.1002/jwmg.21919

Chamberlain MJ. 1999. Ecological relationships among bobcats, coyotes, gray fox, and raccoons and their interactions with wild turkey hens [Ph.D.]. [United States -- Mississippi]: Mississippi State University.

Chamberlain MJ, Leopold BD. 2005. Overlap in Space Use among Bobcats (Lynx rufus), Coyotes (Canis latrans) and Gray Foxes (Urocyon cinereoargenteus). The American Midland Naturalist 153(1):171–179.

Cooper S. 2008. Surveying and habitat modeling for gray foxes in Illinois. Southern Illinois University Carbondale.

Cooper SE, Nielsen CK, McDonald PT. 2012. Landscape factors affecting relative abundance of gray foxes Urocyon cinereoargenteus at large scales in Illinois, USA. Wildlife Biology 18(4):366–373. 10.2981/11-093

Davidson WR, Nettles VF, Hayes LE, Howerth EW, Couvillion CE. 1992. Diseases diagnosed in gray foxes (Urocyon cinereoargenteus) from the southeastern United States. Journal of wildlife diseases 28(1):28–33. 10.7589/0090-3558-28.1.28

Dewitz J. 2023. National Land Cover Database (NLCD) 2021 Products: U.S. Geological Survey data release.

Dore KM, Hansen MF, Klegarth AR, Fichtel C, Koch F, Springer A, Kappeler P, Parga JA, Humle T, Colin C et al. 2020. Review of GPS collar deployments and performance on nonhuman primates. Primates 61(3):373–387. 10.1007/s10329-020-00793-7

Egan ME, Day CC, Katzner TE, Zollner PA. 2021. Relative abundance of coyotes (Canis latrans) influences gray fox (Urocyon cinereoargenteus) occupancy across the eastern United States. Canadian Journal of Zoology 99(2):63–72. 10.1139/cjz-2019-0246

Farias V, Fuller TK, Wayne RK, Sauvajot RM. 2005. Survival and cause-specific mortality of gray foxes (Urocyon cinereoargenteus) in southern California. Journal of Zoology 266(3):249–254. 10.1017/S0952836905006850

Fritzell EK, Haroldson KJ. 1982. Urocyon cinereoargenteus. Mammalian Species (189):1–8. 10.2307/3503957

Hall ER (Eugene R. 1981. The mammals of North America. New York: Wiley.

IDNR. 2005. Illinois Comprehensive Wildlife Conservation Plan & Strategy. Springfield, Illinois, USA: Illinois Department of Natural Resources.

IDNR. 2025. Illinois closes gray fox hunting and trapping season indefinitely.

Larreur MR, Nielsen CK, Lesmeister DB, Bastille-Rousseau G. 2025. The precipitous decline of a gray fox population. Global Ecology and Conservation 58:e03441. 10.1016/j.gecco.2025.e03441

Lesmeister DB, Nielsen CK, Schauber EM, Hellgren EC. 2015. Spatial and temporal structure of a mesocarnivore guild in midwestern north America. Wildlife Monographs 191(1):1–61. 10.1002/wmon.1015

Morris WM, Doak D. 2004. Quantitative Conservation Biology. Sunderland, MA: Sinauer and Associates.

Nicholson WS, Hill EP, Briggs D. 1985. Denning, Pup-Rearing, and Dispersal in the Gray Fox in East-Central Alabama. The Journal of Wildlife Management 49(1):33–37. 10.2307/3801836

Pollock KH, Winterstein SR, Bunck CM, Curtis PD. 1989. Survival Analysis in Telemetry Studies: The Staggered Entry Design. The Journal of Wildlife Management 53(1):7–15. 10.2307/3801296

R Core Team. 2023. _R: A Language and Environment for Statistical Computing_. Rodgers A. 2026. Status, signals and data gaps: The California gray fox. Urban Wildlife Research Project.

Roemer GW, Cypher BL, List R. 2016. Urocyon cinereoargenteus. The IUCN Red List of Threatened Species 2016.

Sikes RS, Gannon WL, the Animal Care and Use Committee of the American Society of Mammalogists. 2011. Guidelines of the American Society of Mammalogists for the use of wild mammals in research. Journal of Mammalogy 92(1):235–253. 10.1644/10-MAMM-F-355.1

Sillero C, Hoffmann M, Macdonald D. 2004. Canids: Foxes, Wolves, Jackals and Dogs. Status Survey and Conservation Action Plan.

Temple DL, Chamberlain MJ, Conner LM. 2010. Spatial Ecology, Survival and Cause-Specific Mortality of Gray Foxes (Urocyon cinereoargenteus) in a Longleaf Pine Ecosystem. The American Midland Naturalist 163(2):413–422.

Therneau T. 2023. A package for survival analysis in R.

Walk J, Ward M, Benson T, Deppe J, Lischka S, Bailey S, Brawn J. 2010. Illinois Birds: A Century of Change.

Weston JL, Brisbin IL Jr. 2003. Demographics of a Protected Population of Gray Foxes (Urocyon cinereoargenteus) in South Carolina. Journal of Mammalogy 84(3):996–1005. 10.1644/BOS-037

Williams BA, Venter O, Allan JR, Atkinson SC, Rehbein JA, Ward M, Marco MD, Grantham HS, Ervin J, Goetz SJ et al. 2020. Change in Terrestrial Human Footprint Drives Continued Loss of Intact Ecosystems. One Earth 3(3):371–382. 10.1016/j.oneear.2020.08.009

